# Dissociable roles for the rTPJ and dmPFC in self-other processing: a HD-tDCS study

**DOI:** 10.1101/306183

**Authors:** A. K. Martin, J. Huang, M. Meinzer

**Affiliations:** The University of Queensland Centre for Clinical Research, Brisbane, Qld, Australia.

## Abstract

**Background:** Social interaction relies on the integration and distinction of self and other. The dorsomedial prefrontal cortex (dmPFC) and the right temporoparietal junction (rTPJ) are two regions consistently associated with social processes. Theories of rTPJ function in social cognition include self-other distinction, self-inhibition, or embodied mental rotation, whereas the dmPFC is associated with a wide range of social functions involving understanding and encoding information pertaining to others. However, to date, no study has provided causal evidence for dissociable roles of the rTPJ and dmPFC in social cognition.

**Method:** 52 healthy young adults were stratified into two studies and received either dmPFC or rTPJ anodal HD-tDCS in a sham-controlled, double-blinded, repeated measures design. Subjects completed a social cognitive battery measuring self-other processing across an implicit and explicit level one (line-of-sight) and level two (mental rotation) visual perspective taking tasks (VPT), as well as self and other encoding effects on episodic memory in order to test the self-reference effect (SRE).

**Results:** Stimulation of the dmPFC selectively enhanced integration of the other perspective into self as indexed by an increase in congruency effect (incongruent-congruent) across both explicit VPT tasks. It also removed the SRE in episodic memory, indexed by increasing the recognition of other-encoded words and reducing the recognition of self-encoded words. Stimulation of the rTPJ resulted in improved inhibition of the self-perspective during level two VPT only, as indexed by a reduction of the congruency effect when taking the other perspective.

**Conclusion:** Our results provide the first causal evidence for dissociable roles of the dmPFC and rTPJ in social cognition. This research supports theories suggesting that rTPJ facilitates embodied mental rotation, whereas the dmPFC integrates information relevant to the other into that of the self.

## Introduction

Integrating and distinguishing between representations related to the self or another person are necessary pre-requisites for higher order social cognition. This meta-representational ability is fundamental to humankind’s ability to empathise with another (i.e feel or understand another’s emotional state) or have a theory of mind (ToM; the ability to understand the beliefs, intentions of another are different from that of one’s own). In this context, the ‘social brain’ is a term used to refer to a network, or set of regions, that are consistently associated with socio-cognitive tasks. Two regions within the social brain are the right temporoparietal junction (rTPJ) and the dorsomedial prefrontal cortex (dmPFC), with these regions implicated in tasks that place demands on self-other processing (Santiesteban, Banissy, Catmur, & Bird, 2012; Schurz et al., 2015; Van Overwalle, 2009; Wittmann et al., 2016).

Specifically, the rTPJ is a highly connected region involved in numerous cognitive processes (Mars et al., 2012), including higher-order social tasks such as ToM (Krall et al., 2016). Competing theories state that the role of the rTPJ in social cognition is either to distinguish between self and other representations (Santiesteban et al., 2012; Schurz, Aichhorn, Martin, & Perner, 2013), shifting to the other representation through inhibition of the self (Payne & Tsakiris, 2017; Soutschek, Ruff, Strombach, Kalenscher, & Tobler, 2016), or more specifically, facilitating embodied rotation and allow the self-perspective to be mentally rotated into an alternate location, including that of other people (Wang, Callaghan, Gooding-Williams, McAllister, & Kessler, 2016). Several theories also exist for the role of the dmPFC in social cognition. Evidence has been put forward for a role in the integration of social information (Brosch, Schiller, Mojdehbakhsh, Uleman, & Phelps, 2013; Ferrari et al., 2016), or a role in integrating information pertaining to self and other in decision-making (Wittmann et al., 2016). However, to date, no study has identified causal and dissociable roles for the dmPFC and rTPJ using tasks able to isolate specific processes relevant to social cognition.

Self-other representations have been measured in a number of ways. Typically, subjects to judge a scene from their own visual perspective or from the hypothetical perspective of an agent or alternate location within a scene. Moreover, visual perspective taking (VPT) can be measured implicitly or explicitly. Here, implicit VPT refers to the automatic tendency to represent another agent’s perspective of a scene without prompting or awareness (Apperly & Butterfill, 2009; Kovacs, Teglas, & Endress, 2010; Ramsey, Hansen, Apperly, & Samson, 2013; Samson, Apperly, Braithwaite, Andrews, & Bodley Scott, 2010). Explicit tasks require the switching from self to other and can be measured on two levels. Level one VPT requires judgements on *if* an object can be seen, whereas level two VPT requires judgement on *how* an object is seen (Michelon & Zacks, 2006). Level one VPT is solvable using “line of sight” judgements whereas Level two VPT is thought to induce a more embodied mental rotation into the other’s perspective and is therefore conceptually closer to ToM (Hamilton, Brindley, & Frith, 2009).

Self-other representations are also important in other cognitive domains. For example, episodic memory is enhanced for items or events that are encoded in relation to the self in comparison to another individual (Symons & Johnson, 1997). SRE in episodic memory represents a task that manipulates self and other processes without relying on mental rotation into another location or the requirement for online control of co-activated self and other representations (Santiesteban et al., 2012). A meta-analysis of self-referential processes using fMRI identified the dmPFC as the key region for other-related processes with less evidence for TPJ involvement (Denny, Kober, Wager, & Ochsner, 2012). This would suggest that the rTPJ is not involved in domain general processing of other-related representations and more has a role in either online control (Santiesteban et al., 2012), inhibition of the self or egocentric bias (Payne & Tsakiris, 2017; Soutschek et al., 2016), or embodied rotation (Wang, Callaghan, Gooding-Williams, McAllister, & Kessler, 2015).

In a previous study, we identified a polarity specific (anodal v cathodal) modulation of dmPFC function when integrating other representations into self-representations across VPT and episodic memory domains (Martin, Dzafic, Ramdave, & Meinzer, 2017). In the current study, we employed the same social cognitive battery to explore the different roles of the dmPFC and the rTPJ. Unlike tasks used in previous studies (e.g. Payne & Tsakiris, 2017; Santiesteban et al., 2012; Wang et al., 2015), this battery allows for other-related processes to be parsed into those related to domain general processing related to another agent, self-inhibition in general, or self-inhibition to facilitate mental rotation and thereby provide causal evidence for the dissociable roles of the dmPFC and rTPJ in self-other processing. We hypothesized dissociable roles for self-other processing, with dmPFC stimulation resulting in increased integration of other into self and rTPJ stimulation increasing the integration of the self into other.

## METHOD

### Participants

Participants: Fifty-two healthy young adults (18–35 yrs) were stratified by sex and assigned to either the sham-controlled dmPFC or rTPJ HD-tDCS double-blinded, crossover studies. Stimulation order was counterbalanced across both stimulation sites. The groups were comparable on neuropsychological functioning, ASQ, anxiety and depression scales. All subjects were tDCS-naïve, not currently taking psychoactive medication or substances, and no history of neurological or psychiatric disorder. All participants provided written consent prior to inclusion in accordance with the Declaration of Helsinki (1991; p.1194), completed a safety screening questionnaire, and were compensated with A$50. The ethics committee of The University of Queensland granted ethical approval.

### Baseline Testing

All participants completed a battery of cognitive tests in order to ensure age-appropriate cognitive status and to ensure site-specific effects of HD-tDCS were not due to underlying cognitive differences between the groups. Tests included the Stroop Test, phonemic and semantic verbal fluency, and the following tests from CogState® computerized test battery (https://cogstate.com): International shopping list, Identification test, One-back, Two-back, Set-shifting test, Continuous paired associates learning test, social-emotional cognition test, and the International shopping list - delayed recall.

Social functioning and recent mental health status were measured using the Autism Spectrum Quotient (ASQ; Baron-Cohen, Wheelwright, Skinner, Martin, & Clubley, 2001) and the Hospital Depression and Anxiety Scale (HADS; Zigmond & Snaith, 1983).

### Transcranial direct current stimulation

The stimulation was administered using a one-channel direct current stimulator (DC-Stimulator Plus®, NeuroConn) and two concentric rubber electrodes (Bortoletto, Rodella, Salvador, Miranda, & Miniussi, 2016; Gbadeyan, Steinhauser, McMahon, & Meinzer, 2016). A small centre electrode (diameter: 2.5 cm) was used at both the dmPFC and rTPJ site. At the dmPFC site, a ring-shaped return electrode (diameter inner/outer: 9.2/11.5cm) was used, whereas a smaller return electrode (diameter inner/outer: 7.5/9.8cm) was used for the rTPJ site due to the position of the right ear (see Figure 2). Safety and focal current current delivery for this montage have been confirmed (Gbadeyan et al., 2016; Martin, Huang, Hunold, & Meinzer, 2017). Electrodes were attached over the target region using an adhesive conductive gel (Weaver Ten20® conductive paste) and held in place with an elastic EEG cap to ensure stable conductive adhesion with the skin. The position of the centre electrode was determined using the 10–20 international EEG system. The dmPFC was located by first identifying FPz and Fz and measuring the distance between the two points. The scalp region overlying the dmPFC was located by locating 15% of the distance from the Fz towards the FPz. This approximated the MNI coordinates (0/54/33), which corresponds to the peak activity in a ToM meta-analysis (Schurz, Radua, Aichhorn, Richlan, & Perner, 2014). The ring electrode was positioned symmetrically around the centre electrode. The rTPJ was located using CP6 of the 10–20 EEG system. In both stimulation conditions, the current was ramped up to 1mA (over 8 seconds). In the “sham” condition the direct current remained at 1 mA for 40 sec before ramping down over 5 seconds. In the active stimulation conditions HD-tDCS was administered for 20 minutes before ramping down. Researchers were blinded to the experimental condition by using the “study-mode” of the DC-stimulator (i.e. a pre-assigned code triggered the respective stimulation conditions). To avoid carryover effects of stimulation, stimulation sessions were conducted with at least 72 hours (3 days) in between. Neurophysiological studies that employed conventional set-ups have confirmed that the effects of single stimulation sessions are short lived (depending on the stimulation parameters approx. 30–60 min). Consequently, typical wash-out times in cross-over studies range from 1–7 days (for reviews see Sarkis, Kaur, & Camprodon, 2014; Stagg & Nitsche, 2011). While HD-tDCS effects on motor evoked potentials may be stronger and slightly delayed compared with conventional tDCS (Kuo et al. 2013), no significant neurophysiological effects were found beyond 120 min after the end of the stimulation for HD-tDCS as well. Therefore, it is safe to assume that three days are sufficient to prevent carry-over effects of the stimulation.

### Visual Perspective Taking Task

The visual perspective task (VPT; Martin et al., 2018) involved three separate tests measuring level one VPT (implicit and explicit) and level two VPT (explicit). All tests involved a street scene with tennis balls, rubbish bins, and either a human avatar or a traffic light directly in front of the gaze of the subject at one of three positions on the street - far, middle, or near (see Figure 1). The traffic light was used as a directional control that should direct attention in a similar manner to the human avatar, but crucially without the ability to hold a perspective of the scene, which was particularly of interest in the implicit VPT task (Apperly & Butterfill, 2009; Samson et al., 2010). The test consisted of 176 trials. In 50% of the trials (n=88) a human avatar was present and in 50% of the trials a traffic light was present. The trials were further separated (50% each, resulting in 44 trials in each condition) by whether the number of balls seen by the subject was congruent or incongruent with that of the human avatar’s view or the number of tennis balls the light would directly hit. This resulted in four conditions; avatar congruent, avatar incongruent, light congruent, light incongruent (see Figure 1). All conditions were balanced for number and location of tennis balls. Each VPT had four counterbalanced versions and subjects were presented with different versions between sessions. All tests were completed in the order; level one implicit, level one explicit, and level two explicit. Subjects were instructed to answer as quickly and as accurately as possible. The stimuli remained on the screen until a response was recorded. A fixation cross was presented for 500ms prior to the stimuli. For the level one and level two VPT, the word “you” or “other” was presented for 750msec prior to the presentation of the scene. Subjects were informed that tennis balls would be hidden from the avatar’s view if a rubbish bin occluded the view or if the tennis ball was behind the avatar. If the traffic light was present, the subjects were instructed to imagine the light radiating out from the traffic light towards the subject and to answer how many tennis balls the light would directly hit. Again, if a bin occluded the light or if the ball was behind the traffic light then the light would not directly hit the ball.

**Figure 1.**
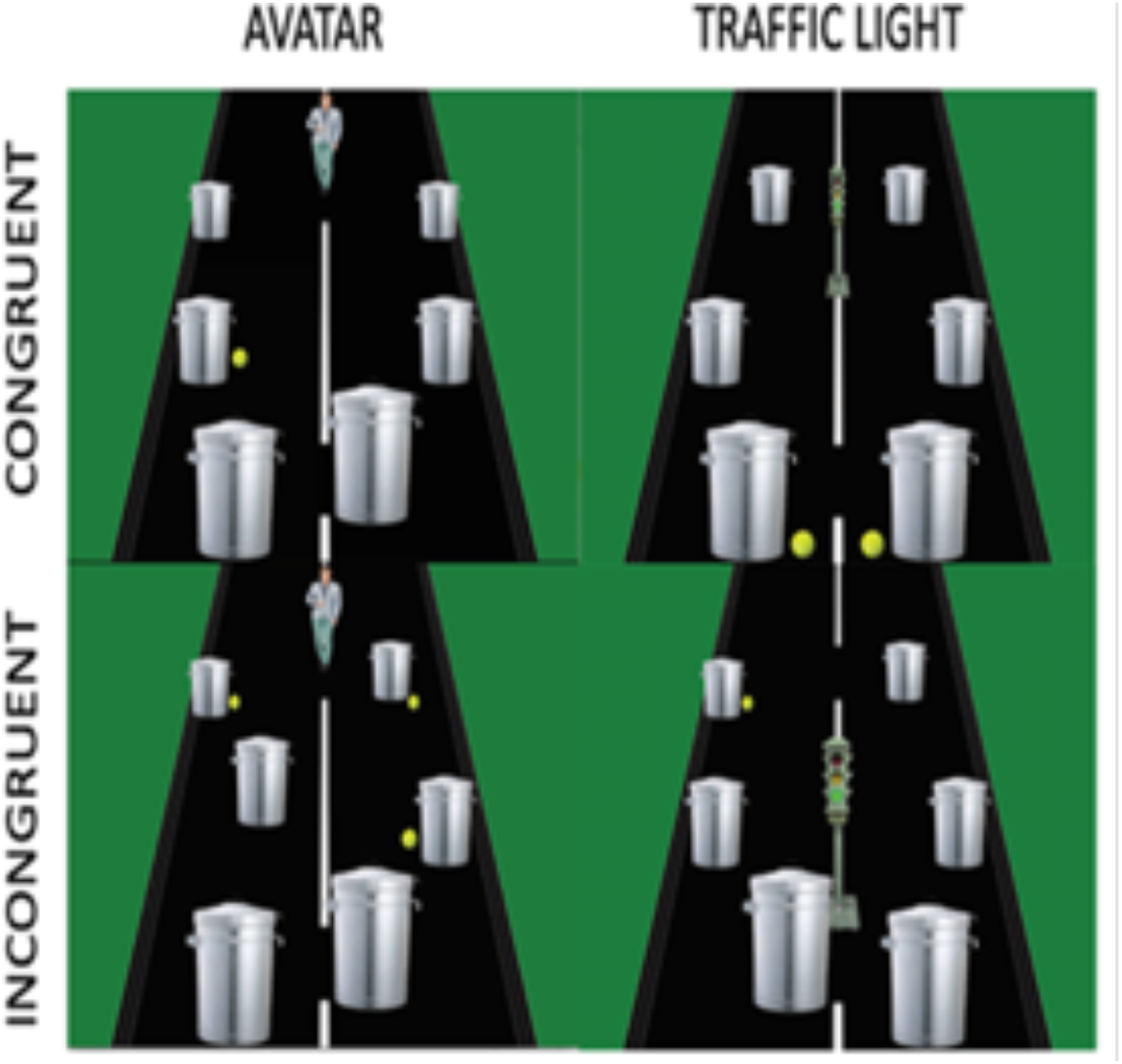
The Visual Perspective Taking (VPT) Task.Examples of congruent and incongruent scenes for both the avatar and the traffic light.

Accuracy and response times are analyzed separately. As the tasks were designed to keep accuracy high, the response time measures are the primary variables of interest. The main outcome of interest was the congruency effect (i.e. the difference between congruent and incongruent trials) and these are plotted in all figures. For the implicit VPT we are interested in agent (avatar v traffic light) specific congruency effects. In both the level one and two explicit visual perspective taking tasks, in line with previous research (Santiesteban, Catmur, Hopkins, Bird, & Heyes, 2014), the congruency effect from the traffic light or avatar was not significantly different for response times, BF_10_= 0.159 and BF_10_= 0.162, respectively, nor accuracy, BF_10_= 0.168 and BF_10_= 0.355, respectively. Therefore, a congruency effect was calculated for both response times and accuracy collapsed across agent. In order to have both RT and accuracy congruency effects in the same direction, congruency effect was calculated as congruent from incongruent for RTs and incongruent from congruent for accuracy. Therefore, higher congruency effect scores for both RTs and accuracy reflect a greater interference from the alternate perspective. For implicit VPT, interference from the avatar and traffic light were calculated separately. It should be noted that wherever stimulation had an effect on congruency effects, these were not reducible to an effect on the incongruent or congruent trials specifically. Instead, stimulation operated at the interaction level between congruent and incongruent trials and either increased or decreased the difference.

### Visual perspective task - Level one implicit

In the first test subjects were instructed to respond as fast and accurately as possible with “how many tennis balls can you see?” The answer was always between one and four with the response buttons clearly marked on the keyboard. The task was considered an implicit test, as subjects were not directed to consider the perspective from the perspective of the avatar in the scene and were only required to answer from the egocentric perspective (see Figure 1).

### Visual perspective task - Level one explicit

In the level one explicit task, participants were required to take either an egocentric perspective or the allocentric perspective from the avatar or light and answer how many tennis balls could be seen. There were four possible responses for each condition, with one to four tennis balls for the egocentric judgements allocentric congruent conditions. In order to maintain four choices for the allocentric incongruent condition, without increasing the number of balls in the scene, scenes with zero balls visible to the avatar/light were included. Therefore, answers in this condition were from zero to three.

### Visual perspective task - Level two explicit

In the level two explicit VPT task, participants were again required to take either an egocentric perspective or the allocentric perspective of the avatar or light. However, this task required making a judgement on “how” the subject or other avatar views the scene, by asking them “whether they/other could see /light would shine on, more balls on the left, right, or equal number on each side of the road?” All conditions had three possible responses.

### Self-referential memory task

Prior to the VPT, participants completed the Reading the Mind in the Eyes Test (RMET; Baron-Cohen, Wheelwright, Hill, Raste, & Plumb, 2001) data published elsewhere (Martin, Huang, et al., 2017). The task requires inferring a person’s mental state solely from the eye region using a four-choice multiple option with a control task requiring the identification of age and sex (Young Male, Young Female, Older Male, Older Female). In order to manipulate the self or other encoding of the memory for the mental attribute, following each choice, the subjects were asked how often they felt that way (self-encoded) or how often they thought Barack Obama felt that way (other-encoded). Prior to the RMET, participants were shown a 5-minute documentary about Barack Obama to ensure familiarization To encourage engagement with the task, subjects were told that their responses would be compared against data collected from people who had worked with Barack Obama.

Following the VPT, participants performed a recognition memory task for the mental attribution words from the RMET. The correct mental attribution words as well as 76 distractor words (38 incorrect choices from the RMET & 38 novel words not previously seen) were presented and subjects answered whether they had seen the mental attribution in the RMET task completed earlier. Responses were; 1= Definitely did, 2= Probably did, 3= Probably not, 4= Definitely not. Scoring was from 2 for a correct confident response through to −2 for a confident response that was incorrect. Words were divided according to whether they had been encoded in relation to the “self” or to the “other” (Barack Obama) and mean confidence scores were calculated.

### Source memory task

If subjects responded that they had seen the mental attribution in the eyes, they were asked a subsequent question “Was it on a male or a female face?” Responses were, 1= Definitely male, 2= Probably male, 3= Probably female, 4= Definitely female. Scoring was identical to the mental attribution memory task. This was considered a source memory, as it was a measure of a contextual memory not directly encoded in relation to the self or other.

For a schematic description of all tasks and stimulation procedures, please see Martin *et al* (2017).

### Adverse Effects and Blinding

Adverse effects were assessed following each stimulation session (Brunoni et al., 2011). Mood before and after stimulation was assessed using the Visual Analogue for Mood Scales (VAMS; Folstein & Luria, 1973). In order to assess blinding, following the final session, subjects were asked to guess which of the two sessions they received the active stimulation.

### Current Modelling

Current modelling (see Figure 2.) was conducted for both the dmPFC and rTPJ stimulation sites (for full details see Martin, Huang, et al., 2017). In brief, modelling of current flow was based on a realistic head model and structural T1-weighted magnetic resonance imaging (MRI) dataset of healthy volunteers. The HD-tDCS simulations were performed using the SimBio software, applying the adjoint approach (Wagner et al., 2014). We obtained the vectorial current density in each finite element generated by HD-tDCS. The current strength was set at 1mA at the central disc electrode and −1mA at the concentric ring electrode. The electrode conductivity was set to 1.4 S/m (Datta, Baker, Bikson, & Fridriksson, 2011).

**Figure 2.**
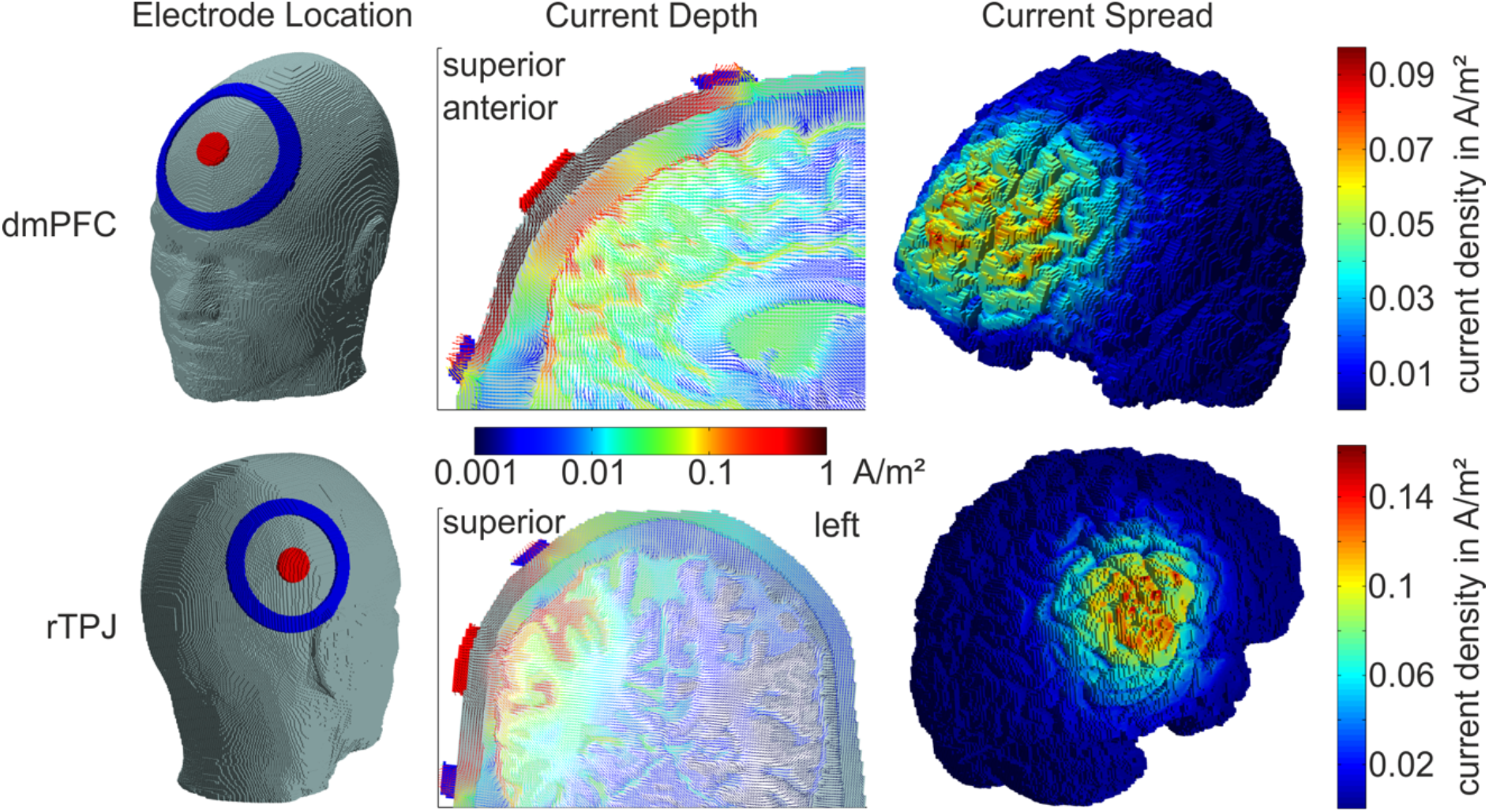
Current modelling for the dmPFC and rTPJ stimulation montages. Image reproduced with permission from Martin *et al* (2017).

### Statistical Analysis

All analyses were computed using JASP version 0.8.6. We applied a Bayesian statistical approach that allowed strength of evidence for both the alternate and null models. A Bayes Factor (BF) quantifies the evidence for a particular model. For example, a BF_10_ of 4 equates to data that is 4 times as likely from the alternate model as from the null model. Evidence for the alternate model is interpreted in a linear scale but for the ease of interpretation we conclude BF_10_ = 1–3 as anecdotal evidence, 3–10 as moderate evidence, >10 as strong evidence. Evidence for the null model follows in the inverse pattern, 0.3–1 anecdotal, 0.1–0.3 moderate, and <0.1 strong evidence (Wagenmakers, Love, et al., 2017). We employed the default priors for all analyses in JASP as recommended (Wagenmakers, Marsman, et al., 2017). The BF_inc_ is the equivalent of the BF_10_ and reports evidence for the inclusion of the main effect or interaction in the model.

Repeated measures analysis of variance (ANOVA) with stimulation location as a between subjects factor (dmPFC and rTPJ) and stimulation type (anodal & sham), perspective (egocentric and allocentric), agent (avatar, light), and congruency (congruent, incongruent) were conducted for the level one and two VPT tasks. The identical analysis was conducted for the implicit VPT minus the perspective condition. For the SRE effect, only stimulation (anodal & sham) and agent (self & other) were included as within-subject factors. All assumptions were met. Individual trials >3 standard deviations from the overall mean were removed from all VPT tasks. Participants who failed to get >50% correct on any condition within the VPT task were removed from that analysis as it was deemed they failed to understand task instructions.

Two subjects from the dmPFC study were removed from the level one VPT analysis as were two subjects from the rTPJ study for level two, for accuracy less than 50%. One subject was removed from the dmPFC level two allocentric analysis as their responses were greater than 4 SDs from the mean and were classified as an outlier. Performance on all VPT and SRE memory measures is provided in Table 1.

**Table 1.** Performance on the Visual Perspective Taking and episodic memory tasks across stimulation type and site. Response times refer to difference between incongruent and congruent trials (msecs) and accuracy is the difference in total correct between congruent and incongruent

|  | dmPFC |  | rTPJ |  |
| --- | --- | --- | --- | --- |
|  | Sham<br>mean (sd) | Anodal<br>mean (sd) | Sham<br>mean (sd) | Anodal<br>mean (sd) |
| *Level two VPT* | N=25 |  | N=24 |  |
| Ego CE RT | 114.87 (139.44) | 193.35 (132.51) | 138.76 (147.85) | 131.12 (190.67) |
| Ego CE Acc | 0.52 (0.92) | 0.42 (1.20) | 0.69 (1.21) | 0.60 (1.22) |
| Allo CE RT | 249.81 (141.56) | 223.16 (163.67) | 244.31 (157.87) | 108.25 (167.26) |
| Allo CE Acc | 1.02 (1.37) | 1.40 (1.33) | 1.15 (1.35) | 1.02 (1.65) |
| *Level one VPT* | N=24 |  | N=25 |  |
| Ego CE RT | 74.31 (79.09) | 122.75 (116.05) | 153.72 (136.35) | 130.87 (128.65) |
| Ego CE Acc | 1.50 (1.98) | 1.48 (1.67) | 2.02 (2.35) | 1.82 (1.98) |
| Allo CE RT | 184.35 (100.88) | 149.38 (145.32) | 208.32 (106.50) | 184.13 (119.40) |
| Allo CE Acc | 0.90 (1.18) | 0.83 (1.04) | 1.32 (1.56) | 1.00 (0.91) |
| *Implicit VPT* | N=26 |  | N=26 |  |
| Avatar RT | 16.29 (18.98) | 12.48 (24.33) | 8.03 (22.72) | 9.97 (25.05) |
| Avatar Acc | 0.00 (1.47) | 0.04 (1.80) | -0.27 (1.59) | 0.27 (1.15) |
| Light RT | -10.38 (20.30) | -12.42 (19.85) | -5.77 (17.82) | -11.74 (20.43) |
| Light Acc | 0.31 (1.74) | -0.50 (1.70) | 0.50 (2.06) | 0.46 (1.68) |
| SRE memory | 0.31 (0.51) | 0.04 (0.41) | 0.00 (0.50) | 0.15 (0.48) |
| SRE source | -0.08 (0.52) | -0.01 (0.43) | 0.05 (0.74) | -0.14 (0.55) |

Ego= Egocentric; Allo= Allocentric; CE= Congruency effect; RT= Response time; Acc= Accuracy; VPT= Visual perspective taking; SRE= Self-reference effect

## Results

### Visual Perspective Taking

#### Level two VPT egocentric

For the congruency effect on RTs, anecdotal evidence was identified for an interaction between Brain Region x Stimulation, BF_inc_= 1.498. Therefore, analyses were conducted for each Brain Region separately. There was moderate evidence for an effect of dmPFC stimulation, BF_10_= 5.803, whereby dmPFC stimulation increased the congruency effect. rTPJ stimulation had no effect, BF_10_= 0.220. Therefore, anodal stimulation to the dmPFC increased the influence or integration of the other perspective with the selfperspective (see Figure 3).

**Figure 3.**
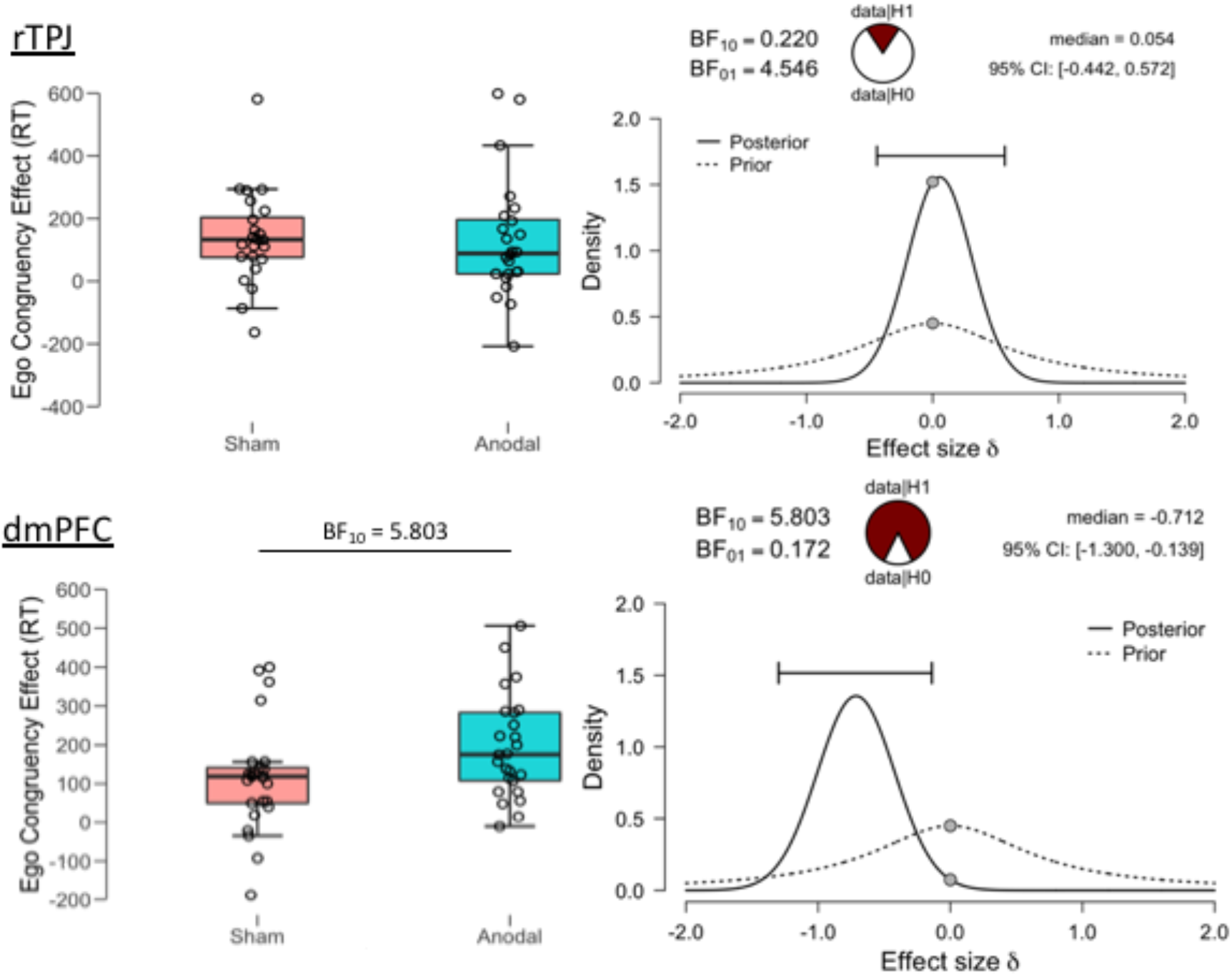
Level two egocentric visual perspective taking. Congruency effect refers to the difference in response time between incongruent and congruent trials. Moderate evidence was provided for an increase in congruency effect after anodal stimulation to the dmPFC. No effects of rTPJ stimulation were identified. Prior and posterior distributions, the median effect size and a 95% credible interval are provided. The pie charts provide a visual representation of the evidence for the null or alternate model.

#### Level two VPT allocentric

For the congruency effect on RTs, anecdotal evidence was identified for an interaction between Brain Region x Stimulation, BF_inc_= 1.383. Therefore, analyses were conducted for each Brain Region separately. There was strong evidence for an effect of rTPJ stimulation, BF_10_= 11.412, such that rTPJ reduced the congruency effect. The null model was supported for dmPFC stimulation, BF_10_= 0.261. Therefore, rTPJ stimulation inhibited the egocentric perspective during a perspective taking task with greater reliance on mental rotation (see Figure 4).

**Figure 4.**
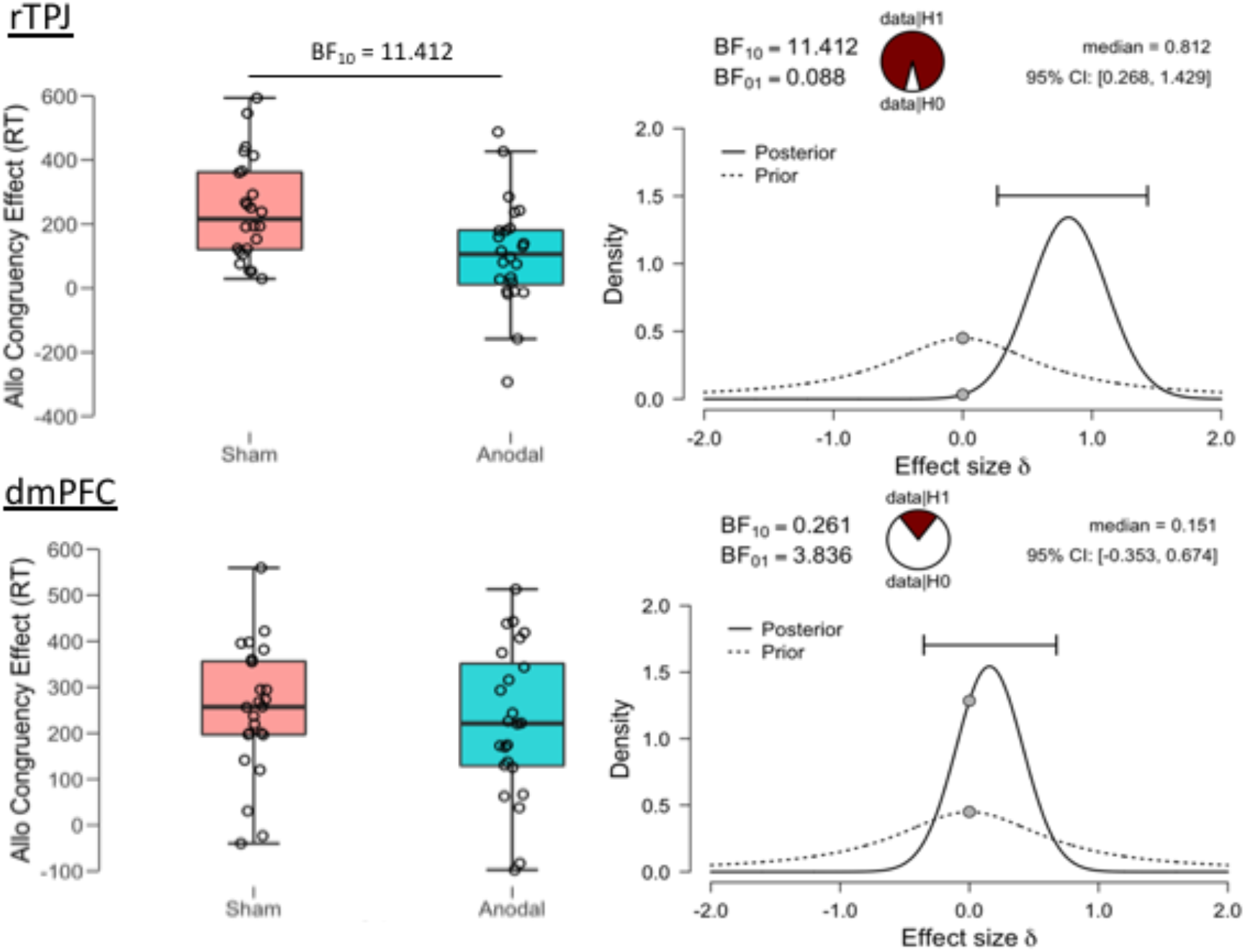
Level two allocentric visual perspective taking. Congruency effect refers to the difference in response time between incongruent and congruent trials. Strong evidence was provided for a reduction in congruency effect after anodal stimulation to the rTPJ. No effects of dmPFC stimulation were identified. Prior and posterior distributions, the median effect size and a 95% credible interval are provided. The pie charts provide a visual representation of the evidence for the null or alternate model.

#### Level one VPT egocentric

For the congruency effect on RTs, anecdotal evidence in support of a Brain Region x Stimulation interaction was identified, BF_inc_ = 1.723. Therefore, simple effects analyses were conducted for the two Brain Regions separately. There was anecdotal evidence in favour for an effect of dmPFC stimulation, BF_10_= 1.012, such that dmPFC stimulation increased the congruency effect. The null model was supported for rTPJ anodal stimulation, BF_10_= 0.334. In a comparable manner to the level two task, dmPFC stimulation increased the integration or influence of the other perspective with the self-perspective (see Figure 5).

**Figure 5.**
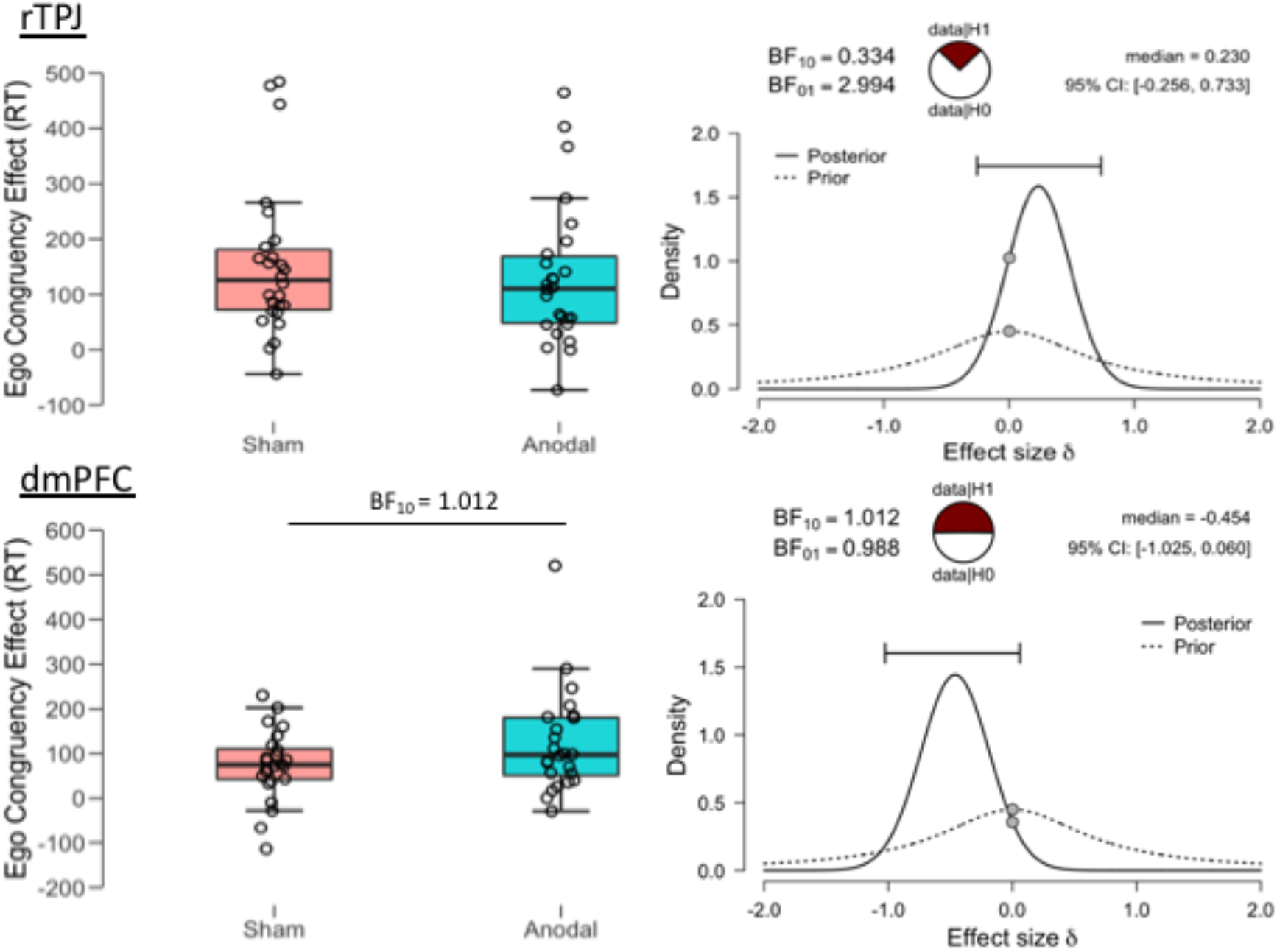
Level one egocentric visual perspective taking. Congruency effect refers to the difference in response time between incongruent and congruent trials. Anecdotal evidence was provided for an increase in congruency effect after anodal stimulation to the dmPFC. No effects of rTPJ stimulation were identified. Prior and posterior distributions, the median effect size and a 95% credible interval are provided. The pie charts provide a visual representation of the evidence for the null or alternate model.

#### Level one VPT allocentric

For congruency effect on RTs, the null model was supported for Stimulation, BF_inc_= 0.654 and for the Brain Region x Stimulation interaction, BF_inc_= 0.302.

### Implicit VPT

An implicit VPT taking effect refers to the automatic tendency to adopt the other’s perspective and is apparent when participant’s are slower to respond to incongruent compared to congruent trials only when an avatar is in the scene and not the traffic light. This is measured in the initial task in which the subjects are only required to answer from their own perspective. An implicit effect was found with strong evidence for an Agent x Congruency interaction, BF_inc_= 182.004, with slower responses when the scene was incongruent with the avatar and surprisingly, the opposite pattern when incongruent with the traffic light. For the congruency effects between both avatar and traffic light, there was no evidence for an effect of Stimulation, BF_inc_= 0.219 nor an interaction between Brain Region x Stimulation, BF_inc_ = 0.200, or Brain Region x Stimulation x Agent, BF_inc_= 0.359.

### VPT Accuracy

There was support for the null model for all stimulation effects on accuracy across all egocentric and allocentric VPT measures and implicit VPT (BF_10_ = 0.178–0.445)

## Self-Reference Effect on Memory

During the baseline sham condition, anecdotal evidence was identified for a self-reference effect for episodic memory (SRE) with greater recognition of words encoded in relation to the self, compared to those encoded in relation to another, BF_10_ = 1.226. The SRE (Self minus Other) was then entered into a RM-ANOVA with stimulation type as a within subject factor and stimulation location as a between subject factor. Moderate evidence was identified for a Brain Region x Stimulation interaction on the SRE, BF_inc_= 4.934. Therefore, paired t-tests were conducted for the effects of stimulation on the SRE for each Brain Region separately. Anecdotal evidence was identified for an effect of dmPFC stimulation, BF_10_= 1.439, such that dmPFC stimulation removed the SRE in episodic memory. After rTPJ stimulation, no effect of stimulation was identified, BF_10_= 0.333 (see Figure 6).

**Figure 6.**
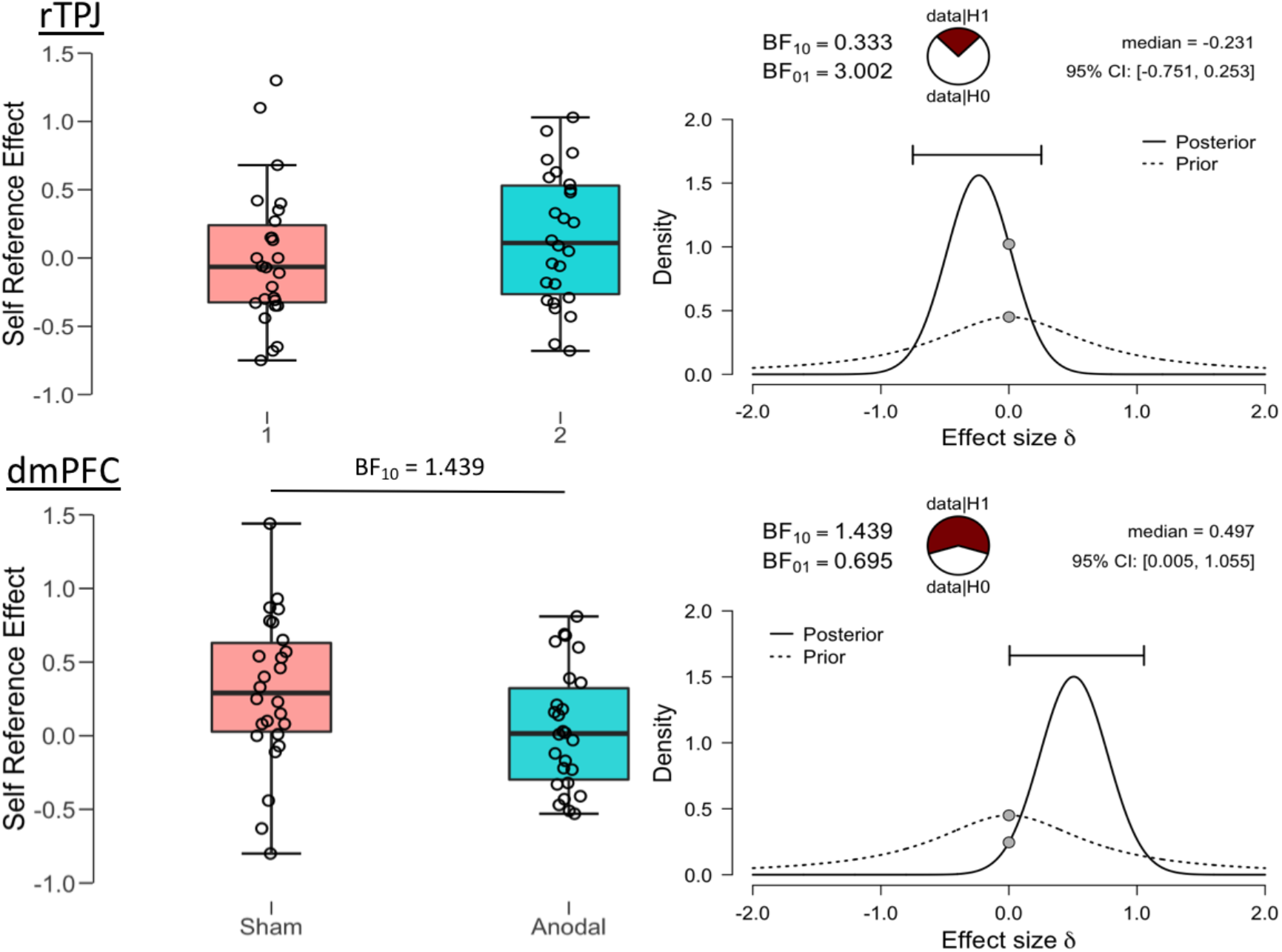
Self-Reference Effect in Episodic Memory. Moderate evidence for an interaction between stimulation sites was identified, BF_10_ = 4.93. Simple effects analyses, demonstrated anecdotal evidence for an effect of anodal tDCS in removing the SRE in episodic memory. rTPJ stimulation had no effect. Prior and posterior distributions, the median effect size and a 95% credible interval are provided. The pie charts provide a visual representation of the evidence for the null or alternate model.

### Source Memory

During the baseline sham condition, no self-reference effect was identified on source memory, BF_10_= 0.154. Stimulation had no effect on source memory, BF_inc_= 0.245 and there was no interaction between Brain Region x Stimulation, BF_inc_= 0.529. Therefore, dmPFC stimulation affected memory only for the items encoded in relation to the self or other and had no effect on the contextual or source memories.

### Baseline Cognition, Adverse Effects, Mood Scales, and Blinding

All participants functioned within age appropriate norms. There was anecdotal evidence for more depressive symptoms, reduced working memory accuracy, and greater number of set-switching errors in the rTPJ group (see Table S1 for details). As the study was a repeated measures design and all subjects were within the normal age-appropriate range, these were not considered in further analyses.

There was no evidence for an effect of Stimulation on adverse effects, BF_inc_ = 0.723 nor was there an interaction between Stimulation x Brain Region, BF_inc_ = 0.505. There was no evidence for an effect of stimulation on increase in negative mood, BF_inc_ = 0.796, or positive mood, BF_inc_ = 0.227 and no interaction between Stimulation x Brain Region for negative mood, BF_inc_ = 0.439 nor positive mood, BF_inc_ = 0.278. Subjects were not able to guess the correct active stimulation session above chance across both studies, BF_10_ = 0.348 (see Table 2).

**Table 2.** Adverse effects and mood scale changes from pre to post stimulation for sham and anodal sessions for both dmPFC and rTPJ studies.

|  | dmPFC |  | rTPJ |  | Stim | StimxRegion |
| --- | --- | --- | --- | --- | --- | --- |
|  | Sham | Anodal | Sham | Anodal | BF<sub>10</sub> | BF<sub>10</sub> |
| VAMS negative | 0.66 (12.04) | 4.11 (6.56) | -0.56 (4.75) | 0.37 (2.90) | 0.80 | 0.44 |
| VAMS positive | -3.47 (28.44) | -4.87 (20.79) | -1.19 (4.49) | -3.15 (7.89) | 0.23 | 0.28 |
| Adverse Effects | 3.35 (2.71) | 4.46 (2.52) | 3.58 (1.96) | 3.77 (2.57) | 0.72 | 0.51 |

## Discussion

This is the first study to identify regionally specific, causal effects of high-definition tDCS on self-other processing. We identified a modulatory effect of dmPFC HD-tDCS on the mergence or integration of other-related processes into the self across cognitive domains. For the right TPJ, we identified a specific effect of inhibiting the self-perspective during allocentric perspective taking with a greater reliance on embodied mental rotation.

Our results provide support for the theory that the rTPJ has a causal role in inhibiting the egocentric perspective during embodied rotation (Wang et al., 2015). As we did not identify a general effect of reducing congruency effects for both self and other processing, our results do not support the theory that rTPJ has a non-specific effect for self-other distinction (Santiesteban et al., 2012). Likewise, we did not find a general self-inhibition effect (Payne & Tsakiris, 2017; Soutschek et al., 2016) as stimulation affected allocentric judgements during level two but not level one VPT. The rTPJ is often associated with ToM or the abilitiy to understand other’s experiences (Krall et al., 2015; Van Overwalle & Baetens, 2009). To date, anodal stimulation to the rTPJ has failed to affect ToM in healthy adults (Martin, Huang, et al., 2017; Santiesteban, Banissy, Catmur, & Bird, 2015) although one study found reduced ToM accuracy after cathodal stimulation of the rTPJ (Mai et al., 2016). As perspective taking, especially the ability to mentally rotate into an allocentric viewpoint, is considered a prerequisite for ToM (Pearson, Ropar, & Hamilton, 2013), the results of the current study, suggest the rTPJ may primarily be involved in lower-order processes relevant for ToM, but not the higher-order ToM ability itself.

The rTPJ is associated with bodily representations (Arzy, Thut, Mohr, Michel, & Blanke, 2006; Blanke & Mohr, 2005; Blanke, Ortigue, Landis, & Seeck, 2002) and specifically implicated in the updated representation of the bodily schema based on proprioceptive and efference-copy information (Branch Coslett, Buxbaum, & Schwoebel, 2008). Therefore, the rTPJ may have a role in imagining the body or mind from a different viewpoint, which may be considered the integration of the self with an external viewpoint. On the contrary, our results suggest the opposite is true for the dmPFC, with a role in the integration of the other into the self, indexed by a greater congruency effect across both explicit VPT tasks. Similarly, it could be interpreted that the removal of the self-reference effect in episodic memory without impairing overall memory after dmPFC stimulation is due to a greater self-encoding of words related to the other.

It has been proposed that social cognition relies on two separate systems, an automatic, implicit system and a conscious, cognitive, explicit system (Apperly & Butterfill, 2009; Frith & Frith, 2008). In the current study, we identified an implicit VPT effect such that incongruent scenes were slower only when an avatar was present and not the traffic light. However, anoal stimulation to the dmPFC or rTPJ had no effect on performance. Both the mPFC and the rTPJ have been implicated in implicit social cognition (Kovacs, Kuhn, Gergely, Csibra, & Brass, 2014) although an alternative account posits that implicit processing occurs in a distinct network of brain regions including the amygdala, basal ganglia, temporal cortex, and the ventral (but not dorsal) portion of the mPFC (Lieberman, 2007). Our results provide causal evidence that the dmPFC and rTPJ are involved exclusively in explicit processes, at least in the domain of visual perspective-taking.

It needs to be noted that level two VPT has been measured using numerous different tasks and a label for a broad range of tasks thought to involve mental rotation (Pearson et al., 2013). Future studies could include additional level two VPT tasks with greater demands on mental rotation to further assess the role of the rTPJ. Although HD-tDCS is more focal than conventional tDCS in the brain regions affected, stimulation effects on underlying brain tissue and connected brain networks remain unknown. For example, several studies that have used conventional tDCS during simultaneous fMRI have demonstrated wide spread modulation of functional networks, primarily in regions that are functionally connected to the stimulation site (Keeser et al., 2011; Meinzer et al., 2012; Meinzer, Lindenberg, Antonenko, Flaisch, & Floel, 2013; Stagg et al., 2013). Similar effects are to be expected for HD-tDCS which could be tested in future studies. Indeed, we have recently demonstrated the feasibility to administer HD-tDCS during fMRI (Gbadeyan et al., 2016). As HD-tDCS avoids current spread to distant brain regions (Bortoletto et al., 2016; Martin, Huang, et al., 2017), such studies could also disentangle stimulation effects due to current spread and direct modulation of neural network nodes functionally connected to the stimulation site. Much work is still required at the basic neurophysiological level to understand how much current reaches the brain and how it alters neuronal function (Huang et al., 2018). However, well controlled studies measuring site and task specificity such as the current study, provide the behavioural evidence to encourage future studies into the plausible explanations of the underlying neural effects of electrical stimulation.

In sum, HD-tDSC to the dmPFC and rTPJ identified dissociable roles in self-other processing. The results support a role for the rTPJ in embodied mental rotation and a role for the dmPFC in the integration of information encoded in relation to the other into that of the self across cognitive domains. We provide causal brain-behaviour evidence explaining how we are able to represent the world from another’s point of view and integrate into our notion of self and thus advancing our knowledge of the social brain.

